# Comparative Aerosol Persistence of Hypermucoid *Klebsiella pneumoniae*

**DOI:** 10.1101/2024.11.07.622510

**Authors:** Nicole Chirichella, Brandon J. Beddingfield, Rachel Redmann, Elizabeth B. Norton, Jay K. Kolls, Chad J. Roy

**Affiliations:** Division of Microbiology, Tulane National Primate Research Center, Covington, LA, USA; Department of Microbiology and Immunology, Tulane School of Medicine, New Orleans, LA, USA; Departments of Pediatrics & Medicine, Center for Translational Research in Infection and Inflammation, Tulane University School of Medicine, New Orleans, LA, USA

## Abstract

*Klebsiella pneumoniae* is becoming an increasing cause for concern due to antibiotic resistance and emerging hypervirulent strains. It results in a wide variety of infections, most commonly pneumonia. Immunocompromised individuals and those needing ventilators are among the most at-risk for infection and systemic spread. It is crucial to further understand the behavior of *K. pneumoniae* to prevent future infections and severe illness. In the past, little research has examined *K. pneumoniae* in the air to determine whether aerosol transmission is a realistic infection pathway. Therefore, we observed survival of K. pneumoniae suspensions using a 10.7 L custom-built rotating chamber, characterizing laboratory and clinical strains in separate aerosol events. Airborne bacterial suspensions were assessed by culture and quantitative polymerase chain reaction (qPCR). The K2 laboratory strain remained viable until yielding no colony forming units (CFU) at 16 hours. K1 clinical strain did not become unculturable until 32 hours. For each strain, qPCR data showed Ct values slightly increased as time in drum also increased. Our data suggests *K. pneumoniae* can survive after multiple hours in the air, indicating aerosol transmission of *K. pneumoniae* is a possible infection pathway.

## Introduction

*Klebsiella pneumoniae* is a gram-negative bacterium causing nosocomial and community-acquired (CA) infections. *K. pneumoniae* is ubiquitous in nature, found in water, soil, plant surfaces, and among the normal microbiota of human and nonhuman mammals ^1,2^. It has also been found in the mouth and on skin. Although classical strains are less likely to harm healthy individuals, those with compromised immune systems are at risk for developing infections leading to meningitis, bacteremia, liver abscesses, and most commonly UTIs and pneumonia ^3^. *K. pneumoniae* causes infection by direct exposure, usually through the respiratory tract. Those most vulnerable to hospital acquired (HA) *K. pneumoniae* infections are patients using ventilators or catheters ^1^. Recently, *K. pneumoniae* has become increasingly resistant to antibiotics, making simple infections concerning, especially with global emergence of hypervirulent (HV) strains ^4^.

Pneumonias can be hospital-acquired (HAP) or community-acquired (CAP), though HAPs are much more prevalent. HAPs present at least 48 hours after hospitalization in patients showing no symptoms of pneumonia prior to admission ^5^. *K. pneumoniae* is among the leading causes of HAPs, along with *Escherichia coli* and *Staphylococcus aureus*. Reportedly, HAPs and ventilator associated pneumonias (VAPs) are the second most frequent cause of nosocomial infections and can have mortality rates of up to 62% ^6^. Those with compromised immunity have a much higher likelihood of presenting with disease caused by classical *K. pneumoniae* and commonly contract infection during hospitalization ^5^. Other risk factors for nosocomial infections include use of corticosteroids, organ transplants, dialysis, chemotherapy, and liver disease ^7,8^. Diabetes, alcohol dependency, and underlying malignancies also substantially increase the risk of HA and CA *K. pneumoniae* infections due to immune suppression. In the future, as diabetes prevalence and HV strains increase, CA K. pneumoniae infections are expected to rise as well ^9^.

The type and severity of *K. pneumoniae* infection is often determined by whether the strain is of classical or HV variety, as well as how the infection was acquired. The rise of HV *K. pneumoniae* strains in Asia and Africa has consequently caused a wider variation in *K. pneumoniae*-associated diseases, many of which are community-acquired ^10^. Unlike classical strains, HV *K. pneumoniae* causes highly dangerous and aggressively-spreading community acquired infections in healthy people ^10^. Morbidity and mortality of pneumonia caused by HV *K. pneumoniae* has also significantly risen. This is in part due to its ability to evade alveolar macrophages and aggressively spread, causing systemic infection ^10,11^. HV strains often grow colonies with higher viscosities, determined by colony size and the string test ^12^.

Though HV *K. pneumoniae* can cause a wide variety of diseases, it is important not to overlook its primary method of transmission: inhalation. Because *K. pneumoniae* easily develops resistance against medications, it is imperative to hinder initial infection along with developing new medications. Therefore, it is crucial to understand the behavior of *K. pneumoniae* in the environment. Considering the most likely infection pathway is through inhalation, understanding modes of transmission in relation to the respiratory tract is among the first steps to lowering infection rates. To date, there has been little research studying the behavior of *K. pneumoniae* aerosol suspensions. Therefore, little is known regarding the abilities and likelihood of *K. pneumoniae* to be transmitted through the air. It is important to determine whether aerosol transmission of *K. pneumoniae* is a realistic pathway for infection in order to assess threat levels, especially for those receiving intensive care. To accomplish this, the following study observed laboratory and clinical strains, K2 and K1 respectively, of *K. pneumoniae* in a rotating chamber to determine length of time each remains viable suspended in air.

## Methods

### Aerosol chamber

We used a customized 10.7 L rotating chamber, or drum, as previously described by Verreault et al. ^13^. All aerosol events took place within a standard Class III Biological Safety Cabinet (GermFree Laboratories, Ormond Beach, FL) measuring 8’×4’×3’, in which the chamber was placed. The chamber included a 45.7 cm long aluminum cylinder, closed by aluminum end caps mounted bilaterally on 5 cm ball bearings. It was supported by pressure-fitted aluminum rods, which remained fixed in position during rotation. A 1/10 hp motor rotated the chamber at a constant speed of 1.0 rpm. Speed was determined by motor capacity and optimized rotation rate calculated using equations found in previous research ^13-15^. A Collison nebulizer (BGI Inc., Waltham, MA) and model 7541 All Glass Impinger (AGI) sampler (Ace Glass Inc., Vineland, NJ) were connected on opposite sides of the chamber. Capsule filters were used to control chamber pressure during sampling and nebulization. Ports were operated manually by ball valves.

### Bacterial Strains

K2 laboratory strain (ATCC, Manassas, VA) and K1 clinical isolate KP-396 were used for this study. Stock bacteria was stored at -80°C until use in aerosol experiments.

### Culture Media and Inoculum Preparation

Each strain was prepared, aerosolized, and sampled separately. Methods were identical for laboratory and clinical strains. From stock bacteria, a lysogeny broth (LB) agar plate was streaked for colony growth.

One colony was added to 100 mL LB medium and incubated at 37°C, 220 rpm overnight. The following day, 100 µl original inoculum was added to 100 mL pre-warmed 37°C LB and incubated again at 37°C, 220 rpm until growth entered log phase, approximately 2-4 hours. Each new inoculum was prepared adding 100 µl of the previous day’s inoculated medium to pre-warmed 100 mL LB.

A growth curve was constructed to identify when growth entered log phase (Fig. S1). OD_600_ values were measured using a Genesys™ 20 Spectrophotometer (Thermo Fisher, Waltham, MA). Growth reached log phase when OD_600_ values measured 0.2A—0.6A. Optimal OD_600_ for this study was defined as 0.4A. Samples of varying OD_600_ values were diluted and plated to identify optimal dilution factor for sample plating.

### Aerosol Projection and Sampling

For consistency, both short-term and long-term aerosol events were carried out using the drum. Each strain was tested separately. Prior to sample nebulization and rotation, the drum was disinfected by running MB-10 (Quip Laboratories Inc., Wilmington, DE) at full flow rate through the aerosol system, followed by flushing filter dilution air for 5 minutes.

Once OD_600_ showed growth had entered log phase, duplicate 1 mL inoculum samples, termed “Pre-Collison,” were aliquoted and stored at -80°C for RNA isolations. 5 mL of the same inoculum was collected to be nebulized into the drum as bacterial aerosol suspensions, deemed “Post-Collison.” These samples were nebulized into the rotating chamber at 18-20 psi for 5 minutes at a generator flow rate of 6.5-7.5 L/min. After nebulization, the remaining volume of Post-Collison liquid was collected and aliquoted in duplicate 1 mL samples and stored at -80°C for RNA isolation.

Each strain’s aerosolization began with a “real time” time point, during which the chamber remained unsealed, and sample was collected as it simultaneously ran through the drum. For all following timepoints, the chamber was sealed and left to rotate at 1.0 rpm for the pre-specified aging period. Both strains were tested until a long-term aging period yielded 0 colonies when samples were plated. K2 laboratory strain was tested at the following timepoints: real time, 1 minute, 10 minutes, 30 minutes, 60 minutes, 2 hours, 4 hours, 8 hours, and 16 hours. Clinical strain rotated similarly with additional 24- and 32-hour aging periods. Each timepoint was done in duplicate.

A 5-minute sampling period followed each rotation, during which the chamber was unsealed and the aerosol contents inside were expelled into 10mL LB supplemented with 40µl antifoam detergent. These samples were labeled “AGI.”

### Dilutions and Plating

Pre-Collison and Post-Collison samples were diluted and plated in duplicate at concentrations of 10^−5^— 10^−8^. AGI samples were diluted and plated from 10^0^—10^−4^. Plates were placed upside-down overnight at 37°C and counted 16-24 hours later. Plates were labeled too many to count (TMC) if more than 300 colonies were present. The number of colonies was then used to determine colony forming units/liter (CFU/L).

### Nucleic Acid purification

Bacterial DNA was isolated from 1 mL undiluted Pre-Collison and AGI samples with the GeneJET DNA Purification Kit (Thermo Fisher, Waltham, MA) according to the manufacturer’s protocol. Procedure was carried out in a DNase-free environment on ice. Isolations yielded 100µl purified DNA in nuclease-free water and were aliquoted into two 50µl samples. Samples were then stored at -80°C until use for qPCR.

### Quantitative polymerase chain reaction

qPCR was done SsoAdvanced Universal SYBR Green Supermix (Bio-Rad Laboratories, Hercules, CA) as per the manufacturers’ protocols. Forward primer (5’-GAT GAA ACG ACC TGA TTG CAT TC -3’) and reverse primer (5’-CCG GGC TGT CGC CAT AAG -3’) were prepared at 100µM concentration according to specification sheets provided (Integrated DNA Technologies, Coralville IA). Primer mix included 1:1:8 FOR:REV:nuclease-free water. qPCR was done in accordance with Bio-Rad’s protocol to yield a total reaction mix volume of 20µl.

## Results

### Colony Morphology

All samples were diluted and plated in duplicate to be counted 16-24 hours later. The K1 clinical strain exhibited a higher mucoviscosity in comparison to the K2 laboratory strain. Clinical *K. pneumoniae* colonies also varied in size and mucoviscosity whereas laboratory *K. pneumoniae* colonies were similar across plates. These observations were consistent across each study group (Figure S2).

### Bacterial viability remains for extended periods of time in suspension

K2 laboratory strain and K1 clinical strain were separately nebulized into a chamber and rotated at different timepoints until duplicate samples yielded 0 colonies when plated. For both strains, Pre-Collison CFU/L remained constant (Figure 1A). K2 laboratory strain CFU/L decreased until plating yielded 0 colonies at 16 hours. K1 clinical strain remained viable longer, reaching 0 colonies at the 32-hour timepoint (Figure 1B).

**Figure 1.**
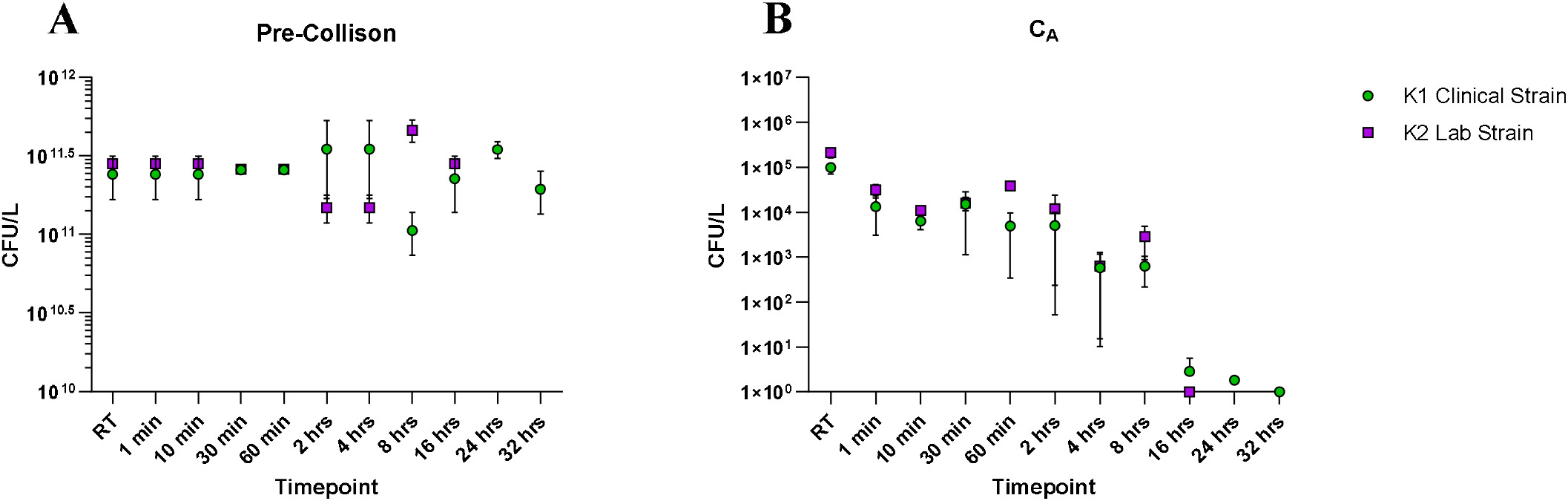
Bacterial titers resulting from continuous suspension of *K. pneumoniae*. Titers from pre-Collison (A) and AGI (B) demonstrating viability in suspension. C_A_: Aerosol concentration.

### Bacterial genome is stable in suspension

Two step RT-qPCR was done to quantify *K. pneumoniae* DNA in each sample by genome copy number. Pre-Collison values (Figure 2A) from each strain remained constant across timepoints. AGI values from each strain showed slight decreases over time (Figure 2B). In relation to each other, laboratory and clinical strains of each group were very similar.

**Figure 2.**
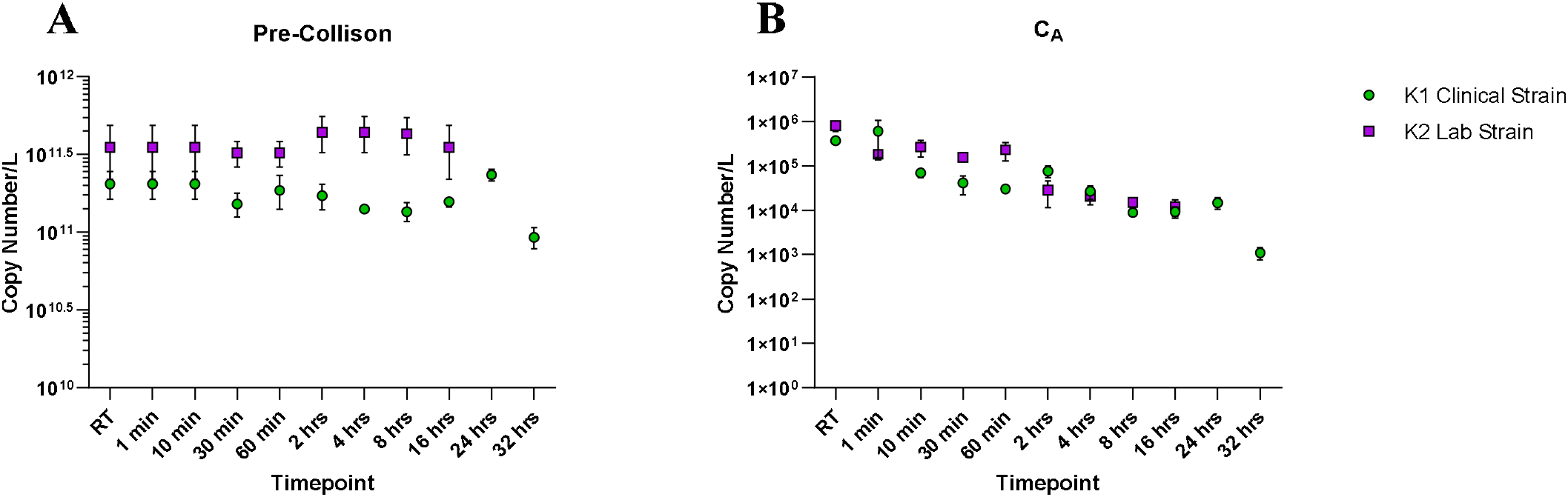
Genome copy numbers resulting from continuous suspension of *K. pneumoniae*. Counts from pre-Collison (A) and AGI (B) demonstrating genome stability in suspension. C_A_: Aerosol concentration.

## Discussion

To date, little research has studied the behavior of *Klebsiella pneumoniae* aerosol suspensions. Pre-Collison CFU values remained constant, while AGI CFU/L values were expected to decrease as rotation time increased. K2 laboratory samples yielded 0 colonies at 16 hours while K1 clinical samples did not become unculturable until 32 hours. Because clinical strain KP-396 was isolated from a hypervirulent case and yielded hypermucoid colonies, it was predicted to remain viable longer than the K2 laboratory strain. This indicates *K. pneumoniae* can survive suspended in air for long periods of time and reinforces the possibility of a transmission route other than direct contact. Still, further research is needed to assess the likelihood of contracting *K. pneumoniae* infections by inhaling aerosol or droplet bacterial suspensions.

The genome of *K. pneumoniae* remains stable over a long period of time in suspension. While pre-Collison values remain constant, AGI genome copies showed a slight increase across time. This minor decrease was constant across groups, with genome of both strains still present after 32 hours, despite the lack of detectable viable bacteria at this time point.

Although the rotating chamber is a well-established method of establishing decay rates of bacterial aerosols, it comes with limitations. For instance, aerosols of larger particle sizes may become deposited onto the walls of the drum, despite centrifugal forces working to prevent this ^16^. Another limitation to utilizing the drum may come from sampling procedures. It is possible the bacteria underwent physical stress during nebulization and sampling processes ^13^. Comparing Pre-Collison and Post-Collison data allows us to determine whether the bacteria experienced significant physical stress due to nebulization. Because the data was very similar, we can be confident this did not occur (data not shown). However, physical stress from AGI sampling is possible, and therefore stability of *K. pneumoniae* aerosols may have been underestimated.

To our knowledge, few studies have been done to determine the longevity of *Klebsiella pneumoniae* in air or on abiotic surfaces. One study observed short-term survival of *K. pneumoniae* aerosols, concluding the bacteria remained living up to 32 minutes ^17^. There have also been reports of *Enterobacteriaceae* species similar to *K. pneumoniae* found in air. One experiment studying workers’ exposure to airborne bacteria reported 35% of bacterial aerosols collected were gram-negative, with the majority belonging to *Enterobacteriaceae* ^18^.

Until recently, few studies have determined the ability of *Klebsiella pneumoniae* to survive long-term suspended in air. Therefore, little was known regarding whether aerosol transmission of *K. pneumoniae* was possible. Our research identified airborne transmission as a realistic pathway for *K. pneumoniae* to cause infection, showing clinical and laboratory strains both survive long-term aerosol events.

Considering past research on similar species of *Enterobacteriaceae*, it may be beneficial for future studies to determine the infectivity of *K. pneumoniae* aerosols *in vivo*. Exposing animal subjects to suspended bacterial aerosols can provide further insight on the likelihood and ability of airborne *K. pneumoniae* to cause infection. This will not only provide insight regarding the aerosols’ ability to infect humans but will also support future methods of prevention.

---

**Figure S1.**
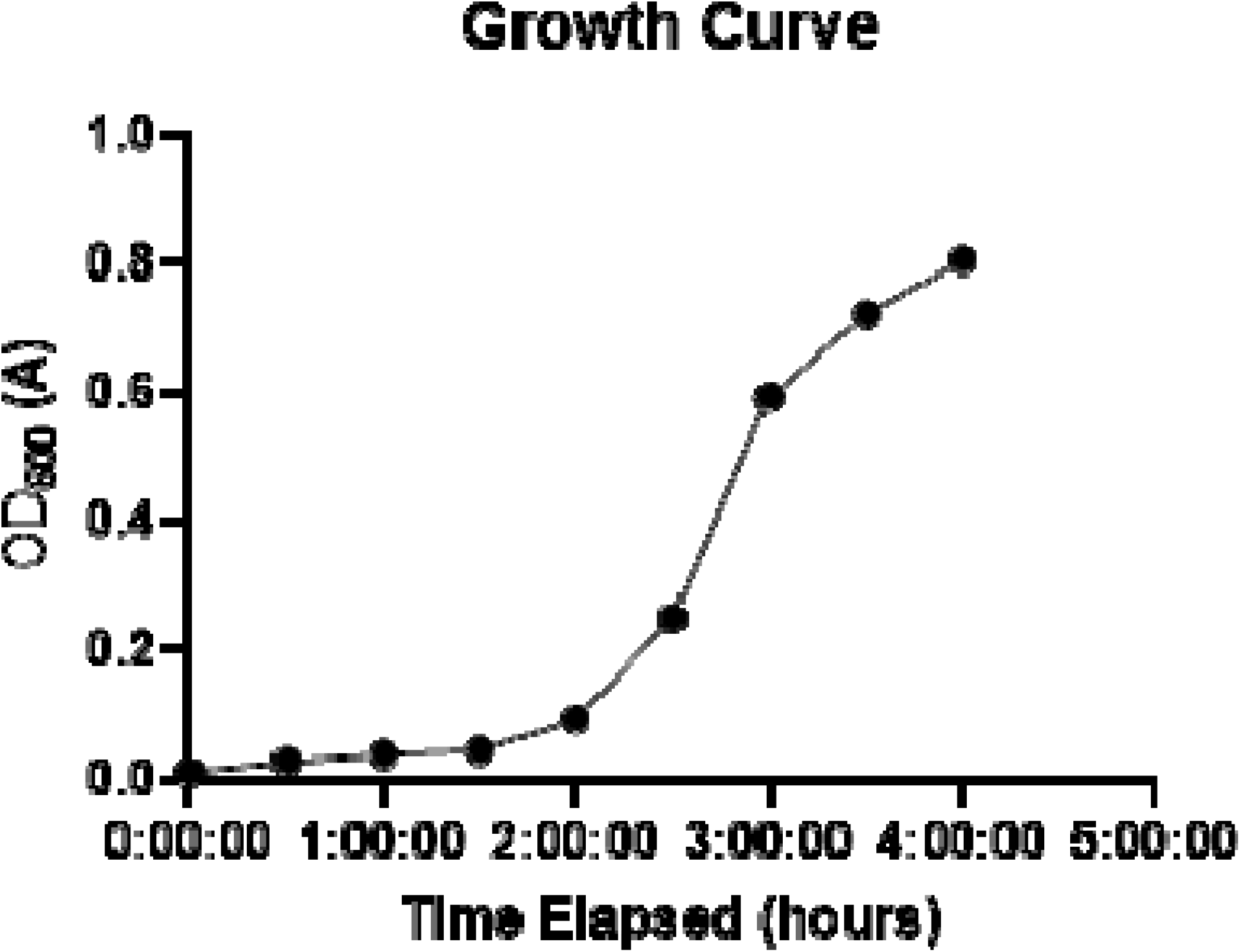
OD600 values were measured every hour to construct the growth curve pictured above. Log phase was identified as 0.2 A—0.6A.

**Figure S2.**
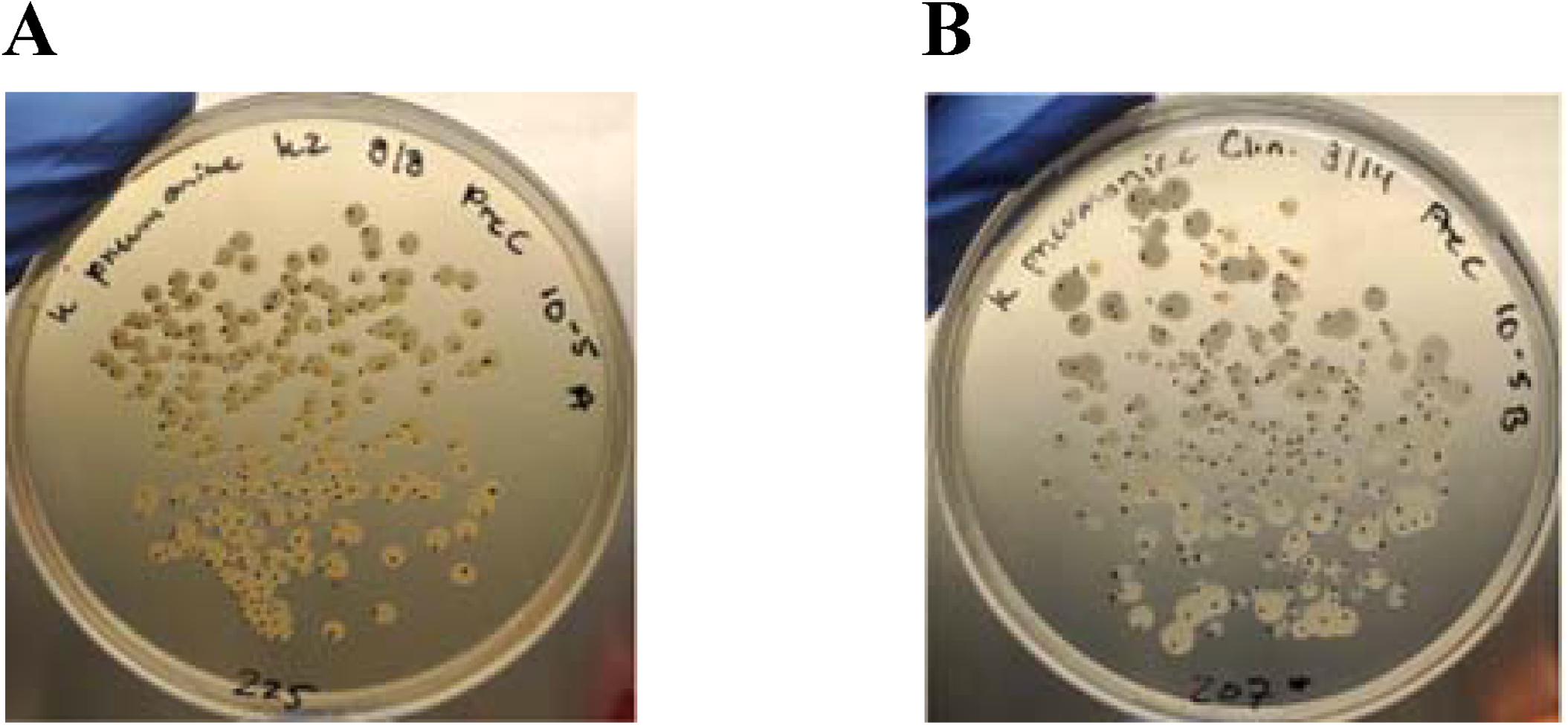
Morphology of plated *K. pneumoniae strains*. Laboratory strain K2 (A) and clinical isolate K1 (B).

